# Endogenous falcarinol-type polyacetylenes in carrots and their putative influence on post-harvest fungal pathogens *Mycocentrospora acerina* and *Botrytis cinerea*

**DOI:** 10.1101/2025.01.08.631919

**Authors:** Wanying He, Frank Dunemann

## Abstract

Falcarinol-type polyacetylenes are a main group of natural bitter compounds synthesized in carrot taproots and putatively contribute to defence mechanisms against soil-borne fungal pathogens. In this study, we analysed the relationships between the constitutive levels of these secondary metabolites and the extent of root infection with the two main carrot storage fungal pathogens *Mycocentrospora acerina* (liquorice rot) and *Botrytis cinerea* (grey mold). Taproots of eight differently colored carrot cultivars exhibiting different levels of the three main falcarinol-type polyacetylenes were inoculated with the two fungi and evaluated for diseased area with a digital image analysis system after 6-weeks incubation in a cold storage facility. After inoculation on cortex tissue and the periderm of taproots, a progressed carrot breeding line and the cultivar Bolero demonstrated the highest tolerance to both *M. acerina* and *B. cinerea*, while Blanche and Senta were the most susceptible cultivars. Concentrations of falcarinol-type polyacetylenes were negatively correlated, though not significantly, with *M. acerina* and *B. cinerea* disease severity. Nevertheless, our study has provided further insights into the naturally occurring variability for the accumulation of falcarinol-type polyacetylenes in cultivated carrots and their putative contributions to resistance against post-harvest fungal pathogens *M. acerina* and *B. cinerea*. To our knowledge, this is the first study about the pathogenicity of *B. cinerea* after inoculation of different carrot cultivars with known root PA levels.

## Introduction

The cultivated carrot (*Daucus carota* subsp. *sativus* Hoffm.) is one of the economically most important vegetables cultivated mainly in temperate regions worldwide (Simon et al. 2019). Carrot taproots accumulate a range of secondary metabolites that can contribute to their bitter off-taste, including falcarinol-type polyacetylenes, laserine-type phenylpropanoids, phenolics like 6-methoxymellein, and distinct terpenes (Kreutzmann et al. 2008; Schmiech et al. 2008). Some of these secondary metabolites were studied for their putative benefits on human health (Cavagnaro et al. 2023), however, their physiological roles in carrots are widely unknown. Plant bitter compounds are believed to play crucial roles in plant defence mechanisms, serving to repel or poison natural enemies, and acting as phytoalexin or phytoanticipin to rapidly enhance resistance in response to abiotic and biotic stresses (Després et al. 2007).

Liquorice rot and grey mold are two main carrot storage fungal diseases, caused by *Mycocentrospora acerina* and *Botrytis cinerea*, respectively. *M. acerina* can infect 80 host plants, including economically important Apiaceae species, such as parsley, celery, carrot and parsnip (Hermansen 1992). *B. cinerea* has over 200 hosts, including grapevine, strawberry, carrot and many other important horticultural crops (Fillinger and Elad 2016). Liquorice rot and grey mold are soil-borne diseases. The main inocula in storage include chlamydospores (*M. acerina*), sclerotia (*B. cinerea*), conidia from infected leaves (*B. cinerea*), and mycelia of both fungi (Williamson et al. 2007, Louarn et al. 2012). These inocula remain in soil debris, adhered to roots, and germinate under optimal conditions. As necrotrophic pathogens, both *M. acerina* and *B. cinerea* infect taproots through wounds caused by machinery injures during post-harvesting processing (Louarn et al. 2012). They secrete toxins to kill host plant cells and utilize enzymes to degrade the cell wall (Le Cam et al. 1994, Amselem et al. 2011). Infected roots exhibit tissue maceration, soft rot, watery parenchyma tissue and in the case of *M. acerina* black lesions with clear margin (Le Cam et al. 1994), while *B. cinerea* infection leads to grey masses of hyphae and conidia (Williamson et al. 2007). These both fungal diseases can cause significant post-harvest losses during storage (Geeson et al. 1988). Currently, no resistant carrot cultivar against *M. acerina* or *B. cinerea* is available.

Polyacetylenes (PA) are a large group of oxylipins derived from fatty acids and widely exist in over 2000 plant species from 24 higher plants families, and they are particularly present in the botanically related families Apiaceae, Araliaceae, and Asteraceae (Konovalov 2014, Dawid et al. 2015). Carrot contains the three major PA falcarinol (FaOH), falcarindiol (FaDOH), and falcarindiol-3-acetate (FaDOAc). They are C_17_-acetylenic compounds with two carbon-carbon triple bonds, two double bonds, and at least one oxygen-containing functional group. PA exhibit diverse bioactivities with human health effects such as antimicrobial, anti-inflammatory, and serotonergic effects (Dawid et al. 2015). Additionally, their cytotoxicity on human cancer cells and potential anticancer effects were established in various preclinical studies (Christensen 2020, Cavagnaro et al. 2023). These PA also exhibit bioactive properties against various plant bacterial and fungal pathogens (Kemp 1978, Hadacek and Greger 2000, Merad et al. 2021). Particularly, FaDOH has been found to exhibit higher toxicity than FaOH against *B. cinerea*, inhibiting the germination or germ tube growth even at low concentrations in *in vitro* assays (Kemp 1978). In the case of *M. acerina*, FaDOH inhibited chlamydospore germination with an IC_50_ (50% inhibition concentration) value of 122 µM and induced structural changes in the cytoplasm of chlamydospores and young hyphal tips at 288 µM (Garrod et al. 1979). *In vitro* tests also showed that FaOH could inhibit the germination of *B. cinerea* spores, and FaOH concentration was induced in root slices treated with *B. cinerea* spores (Harding and Heale 1981). PA distribution in the carrot plant can vary considerably among different plant organs and even within different root segments (Dawid et al. 2015). FaDOH is the most abundant PA in cultivated carrots, with taproot periderm showing the highest total PA levels (Busta et al. 2018). As they accumulated at high levels in root tissues or were being induced under stress conditions, PA are recognized as putative defence metabolites in carrots taproots. Olsson and Svensson (1996) found a weak negative relationship between FaDOH contents in the periderm/pericyclic parenchyma and *M. acerina* infestation and hypothesized that a high endogenous concentration of FaDOH in peripheral tissue could act as a primary barrier against fungal infection of intact roots. As FaDOH was significantly induced in carrot leaves after *Alternaria dauci* infection (Lecomte et al. 2012), and in tomato leaves by *B. cinerea* and *Cladosporium fulvum* (Jeon et al. 2020), this PA is recognized also as a putative phytoalexin.

Carrot pathotests with *B. cinerea* are based on taproot periderm inoculation protocols, while the current inoculation protocols available for *M. acerina* on carrot taproots involve injuring the taproots to the cortex tissues or inoculating cross-section root slices instead of intact taproot structures (Davies et al. 1981, Olsson and Svensson 1996). A periderm-based inoculation method has not yet been described for *M. acerina*. Therefore, the first aim of this study was to explore the relationships between constitutive PA levels in cortex and periderm of different carrot cultivars and the disease severity after inoculation with *M. acerina* on both tissues. Since no published investigation yet exists about the pathogenicity of *B. cinerea* after inoculation of carrot cultivars with different PA levels, the second aim was to investigate the putative influences of differing PA levels in taproot periderm on *B. cinerea* infections.

## Methods

### Carrot material

In this study, eight differently colored carrot cultivars were used, including Blanche (white), Bolero (orange), Gosun (orange), Nerac (orange), Presto (orange), Senta (orange) Yellowstone (yellow) and a progressed orange carrot breeding line (BRL). Carrot seeds were obtained from the JKI working collection (kept by Dr. Thomas Nothnagel) and a Dutch vegetable breeding company (for BRL). Ten seeds were sown and cultivated in 19 cm/30 cm width/height plastic pots in carrot soil in a greenhouse under optimized conditions at 25/20°C day/night and 18 h photoperiod. Two sets of carrots, named set A and set B, were cultivated. Both sets contained the same cultivars, but were sown at different time points and grown in different chambers within the greenhouse. After approximately 200 days of growth, the taproots were harvested. After removing the carrot crowns with leaves using a knife, the taproots were gently washed under running tap water and allowed to air dry before sampling and inoculation.

### Fungal strains, inoculation and disease evaluation

The *M. acerina* strain MA5080 was originally isolated from carrots and kindly provided by a Dutch vegetable breeding company. MA5080 was cultured on potato dextrose agar (PDA) (Carl Roth, Karlsruhe, Germany) at 20°C in the dark. The *B. cinerea* strain BC05.10 used in this study was provided by Prof. Matthias Hahn from the University of Kaiserslautern (Germany). For abundant sporulation, *B. cinerea* was cultured on malt extract agar (Carl Roth) and incubated at room temperature in darkness for 7-10 days.

Carrot set A was used for *M. acerina* (cortex and periderm) and *B. cinerea* inoculation (periderm), while carrot set B was used for a repetition of periderm inoculation with *B. cinerea. M. acerina* chlamydospores inoculum was prepared as described by Day et al. (1972) with modifications. Five fresh fungal plugs from *M. acerina* MA5080 PDA culture were incubated in V8 liquid media (sucrose 30g/L, V8 juice 20 ml/L) in a 500-ml Erlenmeyer flask at 20°C, 110 rpm on a shaker in darkness for two weeks. On the day of inoculation, black clumps of chlamydospores were harvested and blended with sterile water in a stand mixer (Krups Type 240, Germany). The inoculum was filtered through a sieve and adjusted to 1 x 10^5^ chlamydospore chains/ml. For the inoculation of carrot roots, two to three identical wounds were created on each root, spaced 3 cm apart. For each cultivar, we wounded seven to ten roots as replicates shortly before inoculation. To compare the wounding types and the corresponding inoculation performance for *M. acerina*, we implemented two different wound types in carrot set A: the hole wound in the cortex and the scratch wound on periderm. The hole wound was created as described by Davies et al. (1981) with the modification of creating a 3 mm in-depth and 6 mm in-width hole on taproot surface using a 6 mm diameter cork-borer. The scratch wound was created by a 6 mm diameter cork-borer to delimit the wound border and a 1 mm in-depth cross-mark cut with a scalpel. Samples for chemical PA analyses were collected during the wounding process. Cortex samples were tissue plugs from the hole wounds of an individual root, while periderm samples were obtained from peels on the reverse side of a scratch-wounded root using a peeler. Soon after wounding and sampling, an aliquot of 20 µl inoculum was applied to each hole wound and 10 µl to each scratch wound, respectively. Roots inoculated with *M. acerina* were placed in closed plastic trays (length × width × height: 40 cm × 30 cm × 7.5 cm) at 6 °C in the dark for six weeks.

The *B. cinerea* inoculation was conducted as described by Mercier et al. (1993) on carrot surface with modifications. Only periderm inoculations were applied for this fungus. Carrots from set A and set B were wounded with scratch wounds, and periderm samples of set B were collected for chemical analysis as previously described. The *B. cinerea* inoculum was prepared by splashing the culture plate with sterile tap water supplemented with 0.05% Tween 20 and adjusted to 1 x 10^7^ conidia spores/ml. In each wound, 10 µl *B. cinerea* inoculum was applied. To maintain the high humidity and ensure successful infection by *B. cinerea*, the roots were placed on a grid above water-wet paper in closed plastic trays (40 cm × 30 cm × 7.5 cm), and incubated at 4 °C in the dark for six weeks.

The quantitative assessment of diseased areas on taproots was conducted using the LemnaTec digital image analysis system (LemnaTec, Aachen, Germany), which quantified the pixels from an image. Visible light images were processed and analysed with software SAW Bonit (LemnaTec) after calibration for the individual fungal symptoms colours (black for lesion, grey for mycelium) and healthy tissue (healthy taproot colors) (Kathe et al. 2017). For each wounded site, the pixels of respective color classes of diseased tissue were recorded and expressed as the diseased area. In a pilot test (data not shown), area of diseased tissue measured by the LemnaTec system was found to be strongly correlated with the hand-measured diameter of diseased tissue using a ruler (Pearson correlation *r*^2^ = 0.89, *p* < 0.001).

### Analysis of polyacetylenes by HPLC-DAD

The periderm and cortex tissues of carrot set A (used for *M. acerina* inoculation), and the periderm samples of set B (*B. cinerea* inoculation), were kept at −80°C before chemical analysis. Contents of the three falcarinol-type PA compounds FaOH, FaDOH, and FaDOAc were analysed. Sample preparation and metabolites quantification by HPLC-DAD were conducted as previously described (Krähmer et al. 2016).

### Data processing and statistics

Data analysis was performed using R (version 4.1.3). The Shapiro-Wilk test was used to assess data normality. For each individual root, mean diseased area of the wounds were used for analysis. The Kruskal-Wallis test, followed by Dunn’s post hoc test, was conducted on the diseased area at a significance level of *p* < 0.05. The p-values were adjusted using the “Benjamini and Hochberg” method for multiple comparisons. To compare the *M. acerina* diseased area of each cultivar between hole wound (cortex) and scratch wound (periderm), and *B. cinerea* diseased area in periderm between carrot sets A and B, the Mann-Whitney U tests were conducted using wilcox.test function. Due to the non-parametric characteristic of the PA data, we conducted permutational multivariate analysis of variance (PERMANOVA) using the adonis function in the vegan package (Oksanen et al. 2007). PERMANOVA was based on the Bray-Curtis dissimilarity matrix and involved 999 permutations to assess the statistical significance of the differences of polyacetylenes among carrot sets, tissues and cultivars. Spearman rank correlation coefficients between metabolite contents and disease area were computed using the rstatix package (Kassambara 2021).

## Results

### Disease evaluation

Diseased area of inoculated carrots were measured by the LemnaTec digital image analysis system six weeks after inoculation. In the *M. acerina* infection test, carrots of set A were inoculated on periderm and cortex, respectively (Fig. 1a, 1b). Blanche and Senta were the most susceptible cultivars in both tissues, while Bolero, BRL, Nerac and Yellowstone exhibited smaller diseased areas indicating certain levels of tolerance. The breeding line (BRL) showed consistently good tolerance, whereas Bolero appeared to be the best cultivar in both experiments. Notably, as showed by the Mann-Whitney U test, the diseased areas on the periderm were generally smaller than those on the cortex (Suppl. Table S1), especially for Bolero, BRL and Nerac showing significant differences (*p* < 0.05). However, larger diseased areas on periderm were observed for Senta (*p* = 0.041).

**Figure 1.**
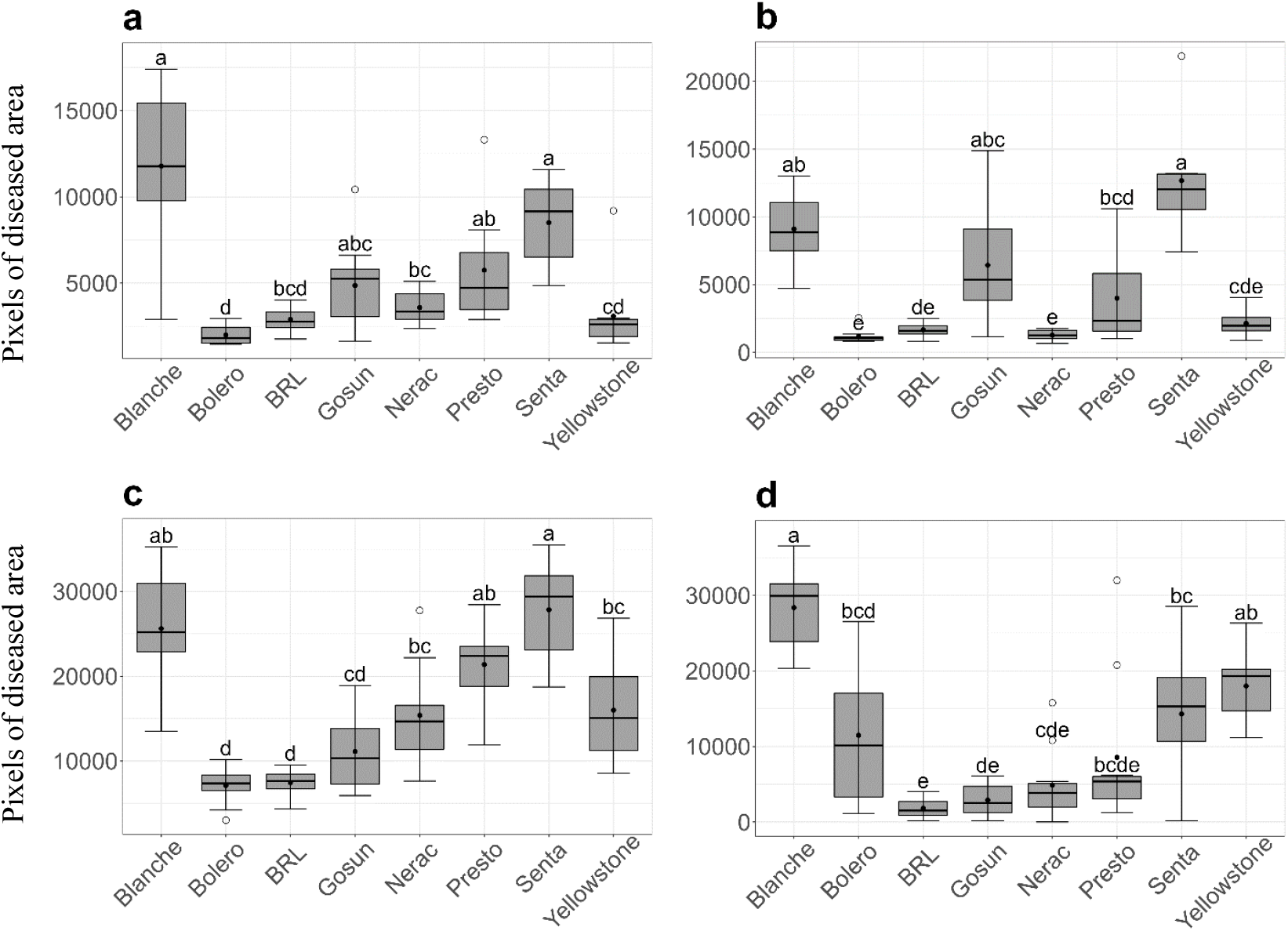
Boxplots of diseased area in different carrot cultivars measured by LemnaTec method in color pixels. **a)** *M. acerina* on carrot set A cortex. **b)** *M. acerina* on set A periderm. **c)** *B. cinerea* on set A periderm. **d)** *B. cinerea* on set B periderm. The solid points indicate the means (n=7 to 10 roots). Letters indicate significant differences among cultivars, determined by the Kruskal-Wallis test and grouped using Dunn’s post hoc test at a significance level of *p* < 0.05. The p-values were adjusted using the “Benjamini and Hochberg” method.

In the *B. cinerea* infections on periderm of set A (Fig. 1c) and set B carrots (Fig. 1d), Blanche (both tests), Senta (set A) and Yellowstone (set B) exhibited the largest diseased areas, indicating the highest susceptibility among the cultivars. The most tolerant cultivars in the set A experiment were Bolero and BRL, while in set B it was BRL followed by Gosun and Nerac. Set A carrots showed generally larger diseased areas and higher susceptibility compared to the taproots of set B (Suppl. Table S2). Compared to the performances in set A, in set B, the diseased areas of BRL, Gosun, Nerac, Presto and Senta were significantly smaller. In contrast, the diseased areas of Blanche, Bolero and Yellowstone were not significantly affected by the different carrot sets.

### Polyacetylene profiling and correlation analysis

Contents of the three PA compounds FaOH, FaDOH, and FaDOAc were quantified using HPLC-DAD in carrot periderm and cortex samples of set A carrots, as well as in periderm samples of set B carrots (Table 1). In the cortex samples of set A, FaOH, FaDOH, and FaDOAc varied between 21 and 662 µg/g DW, 120 and 586 µg/g, and 1.2 and 13.4 µg/g, respectively. The periderm samples generally displayed higher PA levels, ranging from 113 µg/g DW to 1606 µg/g for FaOH and 508 µg/g to 1632 µg/g FaDOH in set A. In periderm, the overall PA levels in set B were generally higher compared to set A, and the levels were significantly influenced by carrot set and cultivar (*p* < 0.001) (Suppl. Table S3). FaDOH was the predominant PA compound in both cortex and periderm tissues, contributing to the overall high level of total PA, while FaDOAc contents showed by far the lowest levels. The highest total PA concentrations were found in both the cortex and periderm of BRL, with concentrations 4 to 8 times higher than those in the low-PA cultivars Blanche and Presto.

**Table 1.**
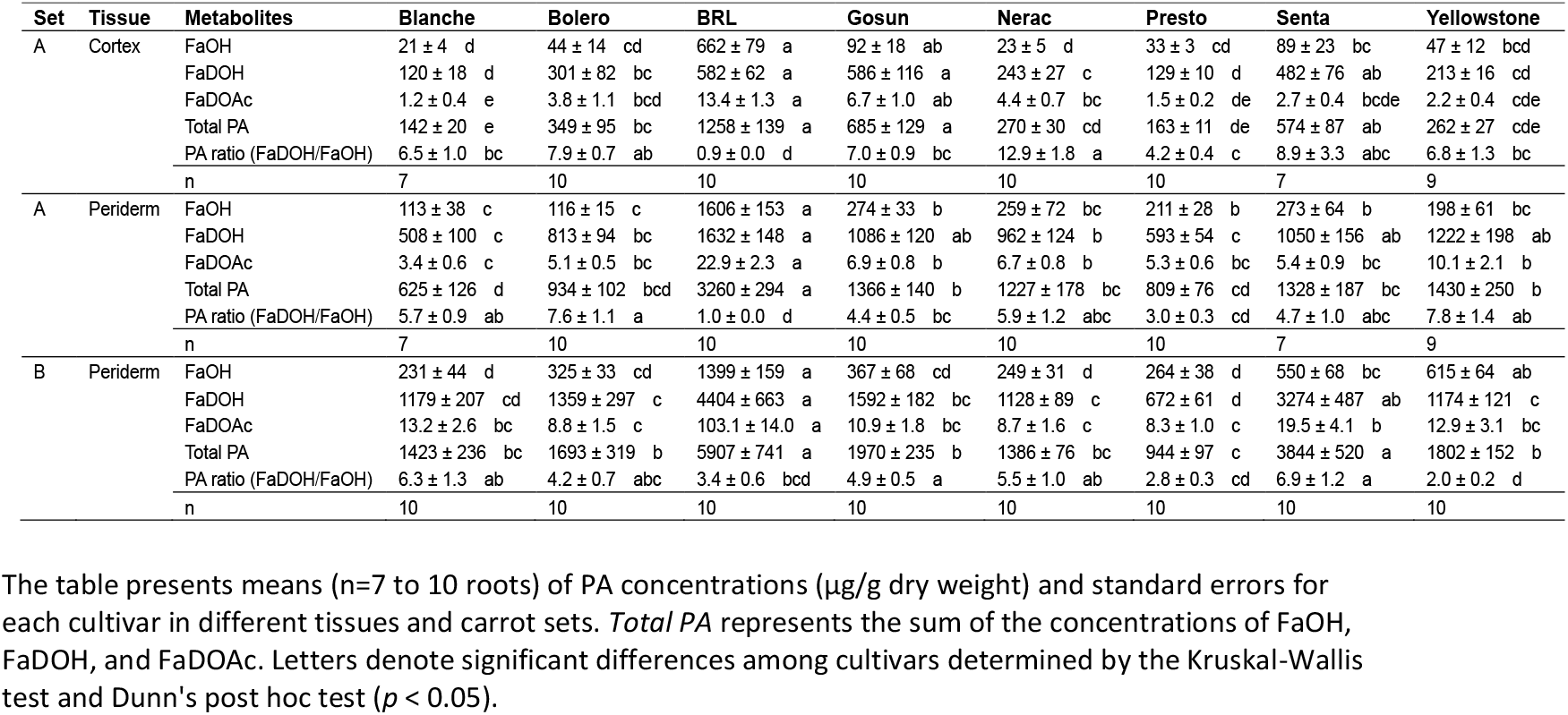
Mean concentrations of falcarinol-type polyacetylenes in carrot cultivars.

Correlation studies were performed to examine the associations between diseased area and PA contents using the following datasets: *M. acerina* inoculation on carrot set A cortex (Table 2a), *M. acerina* inoculation on set A periderm (Table 2b), and *B. cinerea* inoculation on set B periderm (Table 2c). Negative but not significant correlations were observed between diseased area of *M. acerina* and the contents of all PA compounds in cortex (Table 2a). The analysis of diseased area and metabolites contents in periderm also revealed no significant negative correlations for both fungi (Tables 2b, c).

**Table 2.** a) Correlation matrix across *M. acerina* diseased area and PA concentrations in cortex

|  | Diseased<br>area | FaOH | FaDOH | FaDOAc |
| --- | --- | --- | --- | --- |
| FaOH | -0.15 |  |  |  |
| FaDOH | -0.16 | 0.75**** |  |  |
| FaDOAc | -0.21 | 0.62**** | 0.82**** |  |
| Total PA | -0.19 | 0.82**** | 0.98**** | 0.83**** |

**b) Correlation matrix across *M. acerina* diseased area and PA concentrations in periderm**
|  | Diseased<br>area | FaOH | FaDOH | FaDOAc |
| --- | --- | --- | --- | --- |
| FaOH | 0.06 |  |  |  |
| FaDOH | -0.13 | 0.62**** |  |  |
| FaDOAc | -0.23 | 0.60**** | 0.72**** |  |
| Total PA | -0.11 | 0.79**** | 0.96**** | 0.75**** |

**c) Correlation matrix across *B. cinerea* diseased area and PA concentrations in periderm**
|  | Diseased<br>area | FaOH | FaDOH | FaDOAc |
| --- | --- | --- | --- | --- |
| FaOH | -0.11 |  |  |  |
| FaDOH | -0.20 | 0.64**** |  |  |
| FaDOAc | -0.10 | 0.55**** | 0.67**** |  |
| Total PA | -0.17 | 0.78**** | 0.97**** | 0.69**** |

Asterisks indicate significance level: \*\*\*\*p < 0.0001

## Discussion

In our study, PA concentrations were measured by HPLC-DAD in different carrot root tissues used for inoculation with *M. acerina* and *B. cinerea*. The PA quantification showed that all three PA were more concentrated in periderm than in the cortex tissue. This is in agreement with previous findings that these PA are predominantly found in the periderm of carrot taproots, with decreasing concentrations from the outer to inner tissues (Olsson and Svensson 1996, Roman et al. 2011, Busta et al. 2018, Dunemann and Böttcher 2022). Among the PA, FaDOH was found to be the most abundant substance both in periderm and cortex, with levels about 8 to 13-fold higher than FaOH. The abundance of FaDOH and the low concentrations of FaDOAc were also in agreement with results of previous studies (Kjellenberg et al. 2016, Busta et al. 2018). Beside genetic factors several cultivation and environmental factors can influence the content of PA compounds, such as cultivation year, soil conditions, cultivation duration, storage temperature, and processing techniques (Hansen et al. 2003, Dawid et al. 2015, Kjellenberg et al. 2016, Krähmer et al. 2016). In our investigation, the carrots from two distinct sets (sets A and B) exhibited substantial differences in PA concentrations. These differences could potentially be attributed to dissimilar irrigation regimes during the pivotal growth stage, coinciding with taproot expansion, due to the different spatial placement of the two sets within the greenhouse. Additionally, the PA levels varied considerably among cultivars, indicating large genotypic variability for these substance in carrots, as also demonstrated in previous studies (Schmiech et al. 2008, Krähmer et al. 2016, Dunemann and Böttcher 2022).

Our results of the infection experiments with *M. acerina* showed that disease severity was lower following periderm infection compared to cortex infection, while higher PA levels were detected in periderm tissue. This finding is also consistent with the results of Olsson and Svensson (1996). In our study, BRL not only exhibited the highest PA contents but also demonstrated good tolerance to both *M. acerina* and *B. cinerea*. The carrot breeding line BRL is the result of an extensive resistance breeding program targeted to a variety of pathogens. This breeding process also led to the development of high levels of PA, comparable to those observed for example in some *D. carota* ssp. *commutatus* accessions (Dunemann et al. 2022).

Although PA, and among them especially FaDOH, were reported as putative defence compounds in carrots (Harding and Heale 1981, Olsson and Svensson 1996), to our surprise, we were not able to calculate any significant correlation, even not in the case of FaDOH. This finding, however, is in agreement with the study of Olsson and Svensson (1996) who also found no significant relationships between PA contents and resistance against *M. acerina*. Significant correlations were found by these authors only after the elimination of two cultivars, which showed relatively low levels of FaDOH but tolerance to the fungus. Nevertheless, the weak negative correlations found in all three experiments of our study imply that PA might have been indeed contributed to some extent to an improved tolerance. It is likely that the influence of single PA compounds on resistance may have been overruled by other defence-related metabolites which also accumulate in carrot roots, such as the known carrot phytoalexin 6-methoxymellein (6-MM), isocoumarin, terpenes, and laserine-type phenylpropanoids that might be involved in carrot disease response against fungal pathogens (Davies and Lewis 1981, Lecomte et al. 2012, Koutouan et al. 2023, He and Dunemann 2025). For instance, Louarn et al. (2012) reported the induction of 6-MM in *M. acerina-*infected taproots, while no inductions were observed for PA compounds. The cultivar Bolero has shown tolerance against *A. dauci* compared to cultivar Presto (Boedo et al. 2010). Subsequently, Lecomte et al. (2012) observed a higher level of FaDOH and a faster accumulation of 6-MM in Bolero than in Presto during *A. dauci* infection. The induction of 6-MM in carrots, prompted by UV-C (220-280 nm) radiation, resulted in enhanced tolerance against *B. cinerea* (Mercier et al. 1993). In addition, several other reasons might have been responsible for the weak negative correlations between constitutive PA levels and resistance. Apart from chemical defence compounds, carrots have several other resistance mechanisms to prevent pathogen attack, such as a large number of more than 300 putative resistance genes (Boudichevskaia et al. 2022), wound healing effects (Davies and Lewis 1981), and epidermal structure and cell wall properties (Singh et al. 2010). Further investigations on pathogen-induced (elicitated) PA accumulation are needed before an implementation of targeted PA level enhancement into carrot resistance breeding.

## Acknowledgements

We thank Vicky Bartels, Kerstin Maier, and Sabine Struckmeyer (JKI Quedlinburg) for their excellent lab assistance. Astrid Hansen and Dr. Christoph Böttcher (JKI Berlin) are greatly acknowledged for the HPLC analysis. We also thank Dr. Thomas Nothnagel (JKI Quedlinburg) for providing the seeds of the carrot cultivars and Dr. Janine König (JKI Quedlinburg) for suggestions of *B. cinerea* inoculation. We are grateful to Prof. Matthias Hahn (Univ. Kaiserslautern, Germany) who provided the *B. cinerea* strain 05.10, and the Dutch vegetable breeding company which supplied the *M. acerina* strain 5080 and seeds of the progressed breeding line.

## Supplemental Tables

**Table S1.** Mann-Whitney U test for *M. acerina* diseased area between cortex and periderm of set A carrots.

| Cultivar | Cortex median | Periderm median | W value | P value |
| --- | --- | --- | --- | --- |
| Blanche | 11779 | 8854 | 14 | 0.201 |
| Bolero | 1800 | 1037 | 7 | 0.001 |
| Breeding line (BRL) | 2761 | 1575 | 7 | 0.001 |
| Gosun | 5259 | 5365 | 59 | 0.521 |
| Nerac | 3352 | 1255 | 0 | 0.000 |
| Presto | 4727 | 2324 | 29 | 0.121 |
| Senta | 9151 | 12033 | 41 | 0.041 |
| Yellowstone | 2604 | 1963 | 27 | 0.251 |

Results of Mann-Whitney U test for the difference in *M. acerina* diseased area between cortex and periderm tissues of carrot set A for each cultivar. The table presents the median values of the diseased area.

**Table S2.** Mann-Whitney U test for *B. cinerea* diseased area on periderm of carrots of sets A and B.

| Cultivar | Set A median | Set B median | W value | P value |
| --- | --- | --- | --- | --- |
| Blanche | 25176 | 29916 | 32 | 0.505 |
| Bolero | 7354 | 10136 | 39 | 0.427 |
| Breeding line (BRL) | 7633 | 1508 | 100 | 0.000 |
| Gosun | 10307 | 2521 | 99 | 0.000 |
| Nerac | 14674 | 3847 | 92 | 0.002 |
| Presto | 22392 | 5368 | 86 | 0.007 |
| Senta | 29402 | 15288 | 92 | 0.002 |
| Yellowstone | 15074 | 19329 | 38 | 0.385 |

Results of Mann-Whitney U test for the difference in *B. cinerea* diseased area on periderm between carrot set A and set B for each cultivar. The table presents the median values of the diseased area.

**Table S3.** PERMANOVA for PA contents in periderm among set A and set B carrots.

| Source of variation | df | SS | R<sup>2</sup> | Pseudo-F | p |
| --- | --- | --- | --- | --- | --- |
| Cultivar | 7 | 5.69 | 0.45 | 19.5 | 0.001 |
| Set | 1 | 1.07 | 0.08 | 25.7 | 0.001 |
| Residual | 144 | 6 | 0.47 |  |  |

